## Supplementary material for "Divisome minimization shows that FtsZ and SepF can form an active Z-ring, and reveals BraB as a new cell division influencing protein in *Bacillus subtilis*": Saaki SI


- Construction of BMD mutants
- Construction of other mutants
- Table S1 *B. subtilis* strains used in this study
- Table S2 Primers used in this study
- Fig. S1 Growth of Bacillus minimal divisome strains
- Fig. S2 FtsZ and nucleoid localization in BMD27
- Fig. S3 FtsZ and nucleoid localization in *ftsA::erm*
- Fig. S4 Phenotype of the braB deletion mutant
- References

**Construction of BMD mutants**

The minimal divisome mutants were constructed using a marker-free deletion method described by Morimoto et al. (1). A deletion cassette was constructed comprising the upstream and downstream region of the target genes, followed by the IPTG-inducible MazF toxin, a spectinomycin marker for selection and the target gene. This cassette was transformed into the natural competent (transformable) wild type *B. subtilis* 168 cells and selection on spectinomycin containing LB agar plates. After transformation of the donor DNA, the toxin was induced by IPTG, forcing the excision of the deletion cassette by intra-molecular homologous recombination, resulting in deletion of the target gene. After transformation the presence of deletions were checked with PCR. The resulting deletion strain was then used as recipient strain to introduce the next gene deletion.

To create a marker-free *zapA* deletion mutant, the upstream region of *zapA* was amplified with primer pair yshA-DF1/yshA-DR1.2, the downstream region with yshA-DF2.2/yshA-DR2, the MazF-cassette was amplified with yshA-MF/yshA-DR2a from genomic DNA of strain TMO310 (1), and *zapA* was amplified with primer pair yshA-MFa/yshA-MR. The purified PCR products were fused by overlap extension PCR using the flanking primers yshA-DF1 and yshA-MR. The purified final product was transform to competent wild type *B. subtilis* cells resulting in strain *zapA-spec*. MazF toxin induction with IPTG stimulated recombination and removal of the toxin cassette and *zapA*, resulting in strain BMD1.

To create a marker-free *minC* deletion, the upstream region of *minC* was amplified with primer pair minC-DF1/minC-DR1.2, the downstream region with minC-DF2.2/minC-DR2, the MazF-cassette with minC-MF/minC-DR2a from genomic DNA of strain TMO310 (1), and *minC* was amplified with minC-MFa/minC-MR. The purified PCR products were fused by overlap extension PCR using the flanking primers minC-DF1 and minC-MR. The purified final product was transform to competent wild type *B. subtilis* cells resulting in strain *minC-spec*. Chromosomal DNA from this strain was transformed to BMD1, and subsequent MazF toxin induction with IPTG removed the toxin cassette and *minC*, resulting in strain BMD2.

To create a marker-free *ugtP* deletion, the upstream region of *ugtP* was amplified with primer pair ugtP-DF1/ ugtP-DR1.2, the downstream region with ugtP-DF2.2/ugtP-DR2, the MazF-cassette was amplified with ugtP-MF/ugtP-DR2a from genomic DNA of strain TMO310 (1), and *ugtP* was amplified with the primer pair ugtP-MFa/ugtP-MR. The purified PCR products were fused by overlap extension PCR using the flanking primers ugtP-DF1 and ugtP-MR. The purified final product was transform to competent wild type *B. subtilis* cells resulting in strain *ugtP-spec*. Chromosomal DNA from this strain was transformed to BMD2, and subsequent MazF toxin induction with IPTG removed the toxin cassette and *ugtP*, resulting in strain BMD3.

To create a marker-free *minJ* deletion, the upstream region of *minJ* was amplified with primer pair minJ-DF1/minJ-DR1.2, the downstream region with minJ-DF2.2/minJ-DR2, the MazF-cassette was amplified with minJ-MF/minJ-DR2a from genomic DNA of strain TMO310 (1), and *minJ* was amplified with the primer pair minJ-MFa/minJ-MR. The purified PCR products were fused by overlap extension PCR using the flanking primers minJ-DF1 and minJ-MR. The purified final product was transform to competent wild type *B. subtilis* cells resulting in strain *minJ-spec*. Chromosomal DNA from this strain was transformed to BMD3, and subsequent MazF toxin induction with IPTG removed the toxin cassette and *ugtP*, resulting in strain BMD5.

To create a marker-free *ezrA* deletion, the upstream region of *ezrA* was amplified with primer pair ezrA-DF1/ezrA-DR1.2, the downstream pair with ezrA-DF2.2/ezrA-DR2, the MazF-cassette was amplified with ezrA-MF/ezrA-DR2a from genomic DNA of strain TMO310 (1), and *ezrA* was amplified with the primer pair ezrA-MFa/ezrA-MR. The purified PCR products were fused by overlap extension PCR using the flanking primers ezrA-DF1 and ezrA-MR. The purified final product was transform to competent wild type *B. subtilis* cells resulting in strain *ezrA-spec*. Chromosomal DNA from this strain was transformed to BMD5, and subsequent MazF toxin induction with IPTG removed the toxin cassette and *ugtP*, resulting in strain BMD6.

To create a marker-free *spx* deletion, the upstream region of *spx* was amplified with primer pair spx-DF1/spx-DR1.2, the downstream region with spx-DF2.2/spx-DR2, the MazF-cassette was amplified with primerset spx-DR2a/spx-MF from genomic DNA of strain TMO310 (1), and *spx* was amplified with the primers spx-MFa and spx-MR. The purified PCR amplicons were fused by overlap extension PCR with the flanking primers spx-DF1 and spx-MR. The purified final product was transform to competent wild type *B. subtilis* cells resulting in strain *spx-spec*. Chromosomal DNA from this strain was transformed to BMD6, and subsequent MazF toxin induction with IPTG removed the toxin cassette and *ugtP*, resulting in strain BMD7.

To create a marker-free *clpX* deletion, the upstream region of *clpX* was amplified with primer pair clpX-DF1/clpX-DR1.2, the downstream region with clpX-DF2.2/clpX-DR2, the MazF-cassette was amplified with clpX-MF/clpX-DR2a from genomic DNA of strain TMO310 (1), and *ezrA* was amplified with the primer pair clpX-MFa/clpX-MR. The purified PCR products were fused by overlap extension PCR using the flanking primers clpX-DF1 and clpX-MR. The purified final product was transform to competent wild type *B. subtilis* cells resulting in strain *clpX-spec*. Chromosomal DNA from this strain was transformed to BMD7, and subsequent MazF toxin induction with IPTG removed the toxin cassette and *clpX*, resulting in strain BMD9.

To create a marker-free *noc* deletion, the upstream region of *noc* was amplified with primer pair noc-DF1/noc-DR1.2, downstream with noc-DF2.2/noc-DR2, the MazF-cassette was amplified with noc-MF/noc-DR2a from genomic DNA of strain TMO310 (1), and *noc* was amplified with the primer pair noc-MFa/noc-MR. The purified PCR products were fused by overlap extension PCR using the flanking primers noc-DF1 and noc-MR. The purified final product was transform to competent wild type *B. subtilis* cells resulting in strain *noc-spec*. Chromosomal DNA from this strain was transformed to BMD9, and subsequent MazF toxin induction with IPTG removed the toxin cassette and *clpX*, resulting in strain BMD12.

Subsequent removal of *ezrA* from BMD12 was achieved by transforming this strain with chromosomal DNA from strains LH69, which contains the *ezrA::tet* deletion cassette, resulting in BMD14.

The final removal of *ftsA* from BMD14 was achieved by transforming this strain with chromosomal DNA from strains LH75, which contains the *ftsA::erm* deletion cassette, resulting in BMD27. The *ftsA::erm* deletion came from strain YK206 (2), and contains a Pspac promoter that drives expression of *ftsZ*. The promoter does not require IPTG because the *lacI* repressor is not present.

**Construction of other mutants**

To visualize FtsZ using GFP in BMD27, a Pxyl-*gfp-ftsZ* reporter fusion located at the ectopic *amyE* locus was PCR amplified using primer pair TerS350/351 from strain 2020, and DpnI treated to destroy the template DNA. To reduce the chance of deletion mutant reversion and the fact that BMD27 is poorly transformable, the PCR product was first transformed into BMD14, yielding strain TNVS377. Next, genomic DNA of BMD27 was transformed into TNVS377 to delete *ftsA* resulting in strain TNVS385. The PCR product was also transformed to wild type *B. subtilis*, resulting in strain TNVS391. To visualize FtsZ in a *ftsA* deletion strain, TNVS391 was transformed with chromosomal DNA from YK206 (2), resulting in strain TNVS553.

To investigate the localization of BraB, a C-terminal fusion with monomeric super folder GFP was constructed expressed from the ectopic *amyE* locus. To this end *braB* was amplified from genomic DNA using primer pair TerS391/TerS392 and Gibson assembled into pTNV64, which was linearized with primer pair TerS274/TerS368, resulting in plasmid pTNV111. This plasmid was transformed into *B. subtilis* 168 to give strain TNVS298. To remove the native copy of *braB* this strain was transformed with genomic DNA from TNVS34, resulting in strain TNVS308.

To assess the effect of the different spontaneous mutations in the BMD strains, the related genes were deleted. To this end, single deletion strains were obtained from the Bacillus Genomic Stock Centre (BGSC), and their chromosomal DNA transformed to *B. subtilis* 168. The resulting strains are listed in Table S1.

**Table S1:** ***B. subtilis*** **strains used in this study.**

| strain | genotype | reference |
|---|---|---|
| 168 | *trpC2* | (3) |
| TM0310 | *aprE::spec, lacI, Pspac-mazF* | (1) |
| 2020 | *amyE::(Pxyl-gfpmut1-ftsZ spc)* | (4) |
| 1356 | *zapA-yshB::tet* | (5) |
| LH69 | *ΔezrA::tet, amyE::Pxyl-gfp-ftsZ, spec* | Hamoen, unpublished |
| LH75 | *lacA::tet ΔftsA::erm* | Hamoen, unpublished |
| YK206 | *ftsA::erm* $P_{spac}$*-ftsZ* | (2) |
| BKE02340 | *gltP::erm* | (6) |
| BKE00490 | *spoVG::erm* | (6) |
| BKE04730 | *sigB::erm* | (6) |
| BKE06970 | *yesO::erm* | (6) |
| BKE12340 | *uxuA::erm* | (6) |
| BKE13460 | *rsgI::erm* | (6) |
| BKE13470 | *sspD::erm* | (6) |
| BKE13490 | *htpX::erm* | (6) |
| BKE13500 | *ktrD::erm* | (6) |
| BKE13509 | *ykzP::erm* | (6) |
| BKE13510 | *ykzE::erm* | (6) |
| BKE13520 | *ykrP::erm* | (6) |
| BKE13530 | *kinE::erm* | (6) |
| BKE13540 | *ogt::erm* | (6) |
| BKE15050 | *ylbL(ddcP)::erm* | (6) |
| BKE22320 | *ponA::erm* | (6) |
| BKE24770 | *mgsR::erm* | (6) |
| BKE24780 | *yqgY::erm* | (6) |
| BKE24790 | *yqgX::erm* | (6) |
| BKE03220 | *ycgO::erm* | (6) |
| BKE29600 | *braB::erm* | (6) |
| BKE32040 | *yuiF::erm* | (6) |
| BKE36460 | *ywoF::erm* | (6) |
| BKE29530 | *sppA::erm* | (6) |
| BKE21720 | *ypmT::erm* | (6) |
| BKE18670 | *yoaN::erm* | (6) |
| BKE39820 | *htpG::erm* | (6) |
| *minC-spec* | *ΔminC::spec*, $P_{spac}$-*mazF* | this work |
| *zapA-spec* | *ΔzapA::spec*, $P_{spac}$-*mazF* | this work |
| *noc-spec* | *Δnoc::spec*, $P_{spac}$-*mazF* | this work |
| *ugtP-spec* | *ΔugtP::spec*, $P_{spac}$-*mazF* | this work |
| *ezrA-spec* | *ΔezrA::spec*, $P_{spac}$-*mazF* | this work |
| *minJ-spec* | *ΔminJ::spec*, $P_{spac}$-*mazF* | this work |
| *clpX-spec* | *ΔclpX::spec*, $P_{spac}$-*mazF* | this work |
| *spxA-spec* | *ΔspxA::spec*, $P_{spac}$-*mazF* | this work |
| BMD1 | *ΔzapA* | this work |
| BMD2 | *ΔzapA ΔminC* | this work |
| BMD3 | *ΔzapA ΔminC ΔugtP* | this work |
| BMD5 | *ΔzapA ΔminC ΔugtP ΔminJ* | this work |
| BMD6 | *ΔzapA ΔminC ΔugtP ΔminJ ΔezrA* | this work |
| BMD7 | *ΔzapA ΔminC ΔugtP ΔminJ ΔezrA ΔspxA* | this work |
| BMD9 | *ΔzapA ΔminC ΔugtP ΔminJ ΔezrA ΔspxA ΔclpX* | this work |
| BMD12 | *ΔzapA ΔminC ΔugtP ΔminJ ΔspxA ΔclpX Δnoc* | this work |
| BMD14 | *ΔzapA ΔminC ΔugtP ΔminJ ezrA::tet ΔspxA ΔclpX Δnoc* | this work |

| | | |
|---|---|---|
| BMD27 | *ΔzapA ΔminC ΔugtP ΔminJ ezrA::tet ΔspxA ΔclpX Δnoc ftsA::erm* | this work |
| TNVS034 | *braB::erm* (BKE29600 transformed to 168) | this work |
| TNVS083 | *sftA::erm* (BKE29805 transformed to 168) | this work |
| TNVS112 | *ponA::erm* (BKE22320 transformed to 168) | this work |
| TNVS114 | *ycgO::erm* (BKE03220 transformed to 168) | this work |
| TNVS131 | *ylbL(ddcP)::erm* (BKE15050 transformed to 168) | this work |
| TNVS132 | *spoVG::erm* (BKE00490 transformed to 168) | this work |
| TNVS133 | *ypmT::erm* (BKE21720 transformed to 168) | this work |
| TNVS134 | *sppA::erm* (BKE29530 transformed to 168) | this work |
| TNVS135 | *yoaN::erm* (BKE18670 transformed to 168) | this work |
| TNVS193 | *zapA::tet* (1356 transformed to 168) | this work |
| TNVS275 | *uxuA::erm* (BKE12340 transformed to 168) | this work |
| TNVS280 | *glpT::erm* (BKE02340 transformed to 168) | this work |
| TNVS281 | *ftsA::erm* (YK206 transformed to 168) | this work |
| TNVS292 | *braB::erm* (BKE29600 transformed to 168) | this work |
| TNVS298 | *Pxyl-braB-msfGFP* | this work |
| TNVS308 | *braB::erm Pxyl-braB-msfGFP* | this work |
| TNVS377 | *ΔzapA ΔminC ΔugtP ΔminJ ezrA::tet ΔspxA ΔclpX Δnoc* $P_{xyl}$*-gfp-ftsZ* | this work |
| TNVS385 | *ΔzapA ΔminC ΔugtP ΔminJ ezrA::tet ΔspxA ΔclpX Δnoc* $P_{xyl}$*-gfp-ftsZ ftsA::erm* | this work |
| TNVS391 | $P_{xyl}$*-gfp-ftsZ* | this work |
| TNVS401 | *sigB::erm* (BKE04730 transformed to 168) | this work |
| TNVS474 | *yesO::erm* (BKE06970 transformed to 168) | this work |
| TNVS475 | *htpG::erm* (BKE39820 transformed to 168) | this work |
| TNVS476 | *mgsR::erm* (BKE24770 transformed to 168) | this work |
| TNVS477 | *yqgY::erm* (BKE24780 transformed to 168) | this work |
| TNVS478 | *yqgX::erm* (BKE24790 transformed to 168) | this work |
| TNVS479 | *yuiF::erm* (BKE32040 transformed to 168) | this work |
| TNVS515 | *ywoF::erm* (BKE36460 transformed to 168) | this work |
| TNVS516 | *ogt::ermI* (BKE13540 transformed to 168) | this work |
| TNVS517 | *kinE::erm* (BKE13530 transformed to 168) | this work |
| TNVS518 | *ykrP::erm* (BKE13520 transformed to 168) | this work |
| TNVS519 | *ykzE::erm* (BKE13510 transformed to 168) | this work |
| TNVS520 | *ykzP::erm* (BKE13509 transformed to 168) | this work |
| TNVS521 | *ktrD::erm* (BKE13500 transformed to 168) | this work |
| TNVS522 | *htpX::erm* (BKE13490 transformed to 168) | this work |
| TNVS523 | *sspD::erm* (BKE13470 transformed to 168) | this work |
| TNVS524 | *rsgI::erm* (BKE13460 transformed to 168) | this work |
| TNVS553 | *ftsA::erm amyE::(*$P_{xyl}$*-gfpmut1-ftsZ spc)* | this work |

**Table S2: primers used in this study.**

| Primer | Sequence 5'-3' * |
|---|---|
| TerS274 | CATCCTAGGAATCTCCTTTCTAGA |
| TerS350 | CACCGCCGACATTCGCGTGGCTCCA |
| TerS391 | AGAAAGGAGATTCCTAGGatgAAACACTCACTGCCTGTCA |
| TerS392 | CCTGAGCCGCTTCCTGAGCCACTTATTTCATTAAGCTGTTTGGA |
| TerS351 | GCATCAGGGCTGCGGCATCCGGA |
| TerS368 | GGCTCAGGAAGCGGCTCAGGATCCAAAGGAGAAGAACTTTTCACTGGAGT |
| yshA-DF1 | AAGCCCTGACAAGTACGGTG |
| yshA-DF2.2 | gacgctccagGCCGTCAGACAACGTTTCTC |
| yshA-DR1.2 | gtctgacggcCTGGAGCGTCAGCTTAAAGA |
| yshA-DR2 | CTGATTGGGTAggatccgcgCCTTGAAACAAGACCTTACC |
| yshA-DR2a | GGTAAGGTCTTGTTTCAAGGcgcggatccTACCCAATCAG |
| yshA-MF | GTCAAATTACAAGAGAAATGTGcgacagcggaattgactc |
| yshA-MFa | gagtcaattccgctgtcgCACATTTCTCTTGTAATTTGAC |
| yshA-MR | AAACAACCGTTGACATTTACGG |
| minC-DF1 | CCAGCAGTCCTATAATACGG |
| minC-DF2.2 | ttaaatgagcGGTCTTCACAATATTCACCTC |
| minC-DR1.2 | tgtgaagaccGCTCATTTAAGACCTGATCTA |
| minC-DR2 | CTGATTGGGTAggatccgcGCTATTTGGCTTCAAAAGCG |
| minC-DR2a | CGCTTTTGAAGCCAAATAGCgcggatccTACCCAATCAG |
| minC-MF | TCATGTGTTAAATCGCTCCCcgacagcggaattgactc |
| minC-MFa | gagtcaattccgctgtcgGGGAGCGATTTAACACATGA |
| minC-MR | TGGACTAACATTGCATCTGG |
| ugtP-DF1 | TGGCGATTGAAATCCAGATG |
| ugtP-DF2.2 | gcactttggcGGTATTCAAGTAAATTCACCTC |
| ugtP-DR1.2 | cttgaataccGCCAAAGTGCTATCGTAATG |
| ugtP-DR2 | CTGATTGGGTAggatccgcgTGCAAAAGGCAGCAACGAGC |
| ugtP-DR2a | GCTCGTTGCTGCCTTTTGCAcgcggatccTACCCAATCAG |
| ugtP-MF | GTGCTTCTGATCATGGCAGGcgacagcggaattgactc |
| ugtP-MFa | gagtcaattccgctgtcgCCTGCCATGATCAGAAGCAC |
| ugtP-MR | CCAAGAGTCAAATCCGATTG |
| minJ-DF1 | CGATCTTCATGTCAGCCAGC |
| minJ-DF2.2 | cgtcttcacgAACAGACACTATCTCTCACC |
| minJ-DR1.2 | agtgtctgttCGTGAAGACGAAGCAGTCGC |
| minJ-DR2 | CTGATTGGGTAggatccgcGGTCAATTACACCAATGTCG |
| minJ-DR2a | CGACATTGGTGTAATTGACCgcggatccTACCCAATCAG |
| minJ-MF | GAAGCGCTTCAGCATAACCGcgacagcggaattgactc |
| minJ-MFa | gagtcaattccgctgtcgCGGTTATGCTGAAGCGCTTC |
| minJ-MR | CGTTCTGCGCATGTGAGAAC |
| ezrA-DF1 | GCAGCGATTACAGTCGCATG |
| ezrA-DF2.2 | atatgtcagcAATGACAAACTCCATAATGAGC |
| ezrA-DR1.2 | gtttgtcattGCTGACATATCCGCTTAGATA |
| ezrA-DR2 | CTGATTGGGTAggatccgcgAAGGCTTCTTCTGCATATTGTC |
| ezrA-DR2a | GACAATATGCAGAAGAAGCCTTcgcggatccTACCCAATCAG |
| ezrA-MF | GACTAGCAACATTCCAGGCAcgacagcggaattgactc |
| ezrA-MFa | gagtcaattccgctgtcgTGCCTGGAATGTTGCTAGTC |
| ezrA-MR | AATCCTGCAGCTCATTGACG |
| clpX-DF1 | GCTTCTTCCAACACAAGCCG |
| clpX-DF2.2 | cctcagtgccGAACGAGCATTTTAATTGTCCT |
| clpX-DR1.2 | atgctcgttcGGCACTGAGGTAAGCCAAGA |
| clpX-DR2 | CTGATTGGGTAggatccgcgAAGCCATCGGTTCTACTGAC |
| clpX-DR2a | GTCAGTAGAACCGATGGCTTcgcggatccTACCCAATCAG |
| clpX-MF | CTTCCAGAAGATTTGCTCCGcgacagcggaattgactc |
| clpX-MFa | gagtcaattccgctgtcgCGGAGCAAATCTTCTGGAAG |

| | |
|---|---|
| clpX-MR | AGCGCATTAACTCCAACAGC |
| noc-DF1 | AACGAAGAGAGATGTACCCG |
| noc-DF2.2 | tgcgaatcgtGAATGAATGCTTCATGTACCTA |
| noc-DR1.2 | gcattcattcACGATTCGCATACCAAAATAG |
| noc-DR2 | CTGATTGGGTAggatccgcgAAAAGCCGCATCAGCTGAAG |
| noc-DR2a | CTTCAGCTGATGCGGCTTTTcgcggatccTACCCAATCAG |
| noc-MF | GAACACGATTCGCCAGTCACcgacagcggaattgactc |
| noc-MFa | gagtcaattccgctgtcgGTGACTGGCGAATCGTGTTC |
| noc-MR | GAAGAAGGCCAATACGAACTC |
| spx-DF1 | CATAAAGATGAAGGCAAACATC |
| spx-DF2.2 | gctaattgagGGCAAACTAATAGATCGTATC |
| spx-DR1.2 | ttagtttgccCTCAATTAGCTTAACTGATCGAA |
| spx-DR2 | CTGATTGGGTAggatccgcGGACAACGTCATCGCAGTTG |
| spx-DR2a | CAACTGCGATGACGTTGTCCgcggatccTACCCAATCAG |
| spx-MF | GAAAAAAATGTGTCAATCCAGCgacagcggaattgactc |
| spx-MFa | gagtcaattccgctgtcGCTGGATTGACACATTTTTTTC |
| spx-MR | CTTCTCTTAATTGGAAAGAGCG |

* matching sequences in capital letters.

**Fig. S1.**

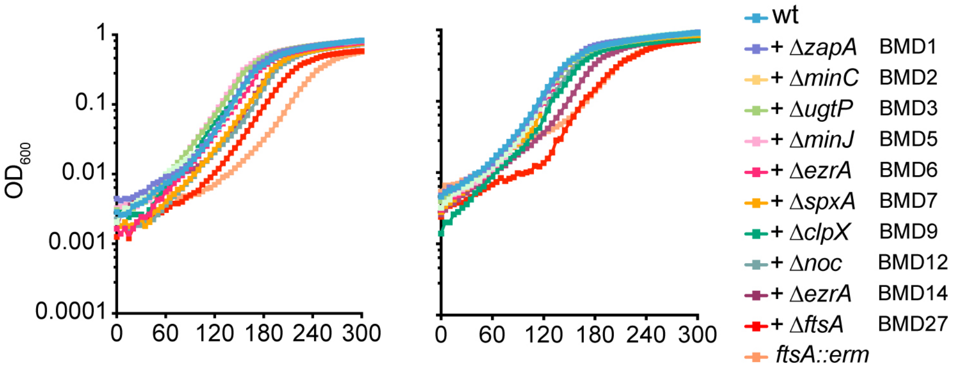


**Fig. S1. Growth of Bacillus minimal divisome strain.**

Growth curves of BMD strains grown in microtiter plates at 37 $^{o}$C in LB medium supplemented with 1 % glucose and 10 mM $MgSO_4$. Biological replicates related to Fig. 2B in main text.

**Fig. S2.**

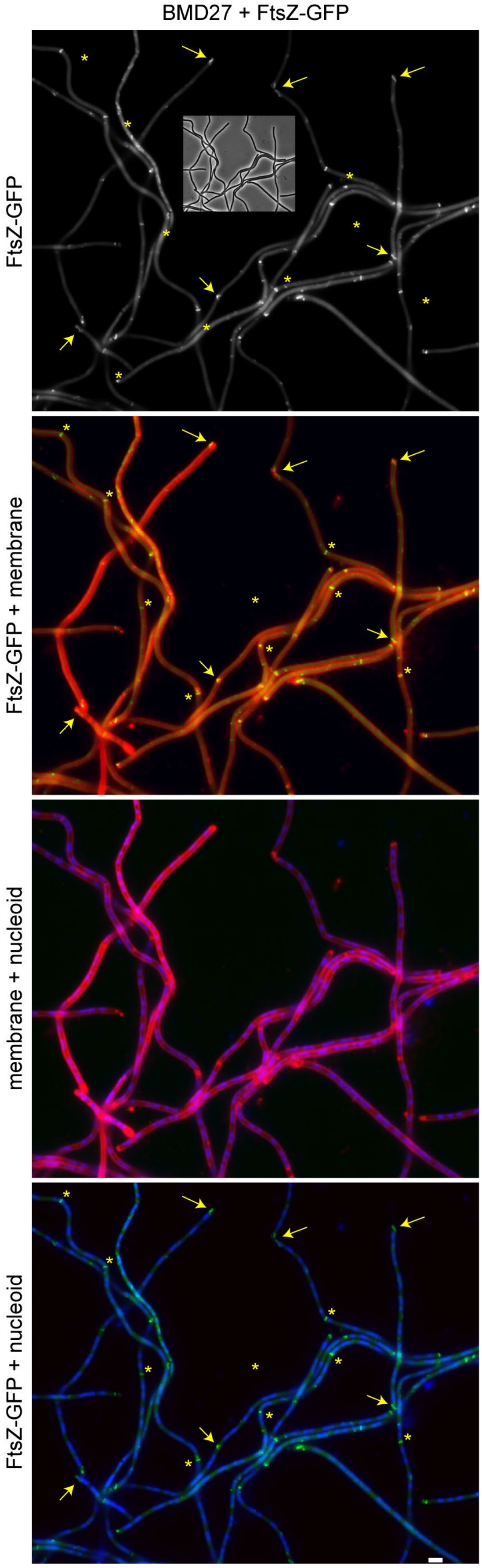


**Fig. S2. FtsZ and nucleoid localization in BMD27.**

Fluorescence microscopy images of exponential growing BMD27 cells expressing an inducible ectopic FtsZ-GFP reporter protein (strain TNVS385). The phase contrast image is shown as inset in the FtsZ-GFP (top) panel. Membrane (red) and nucleoids (blue) were stained with FM5-95 and DAPI, respectively (2$^{de}$ and 3$^{de}$ panel, respectively). Asterisks indicate FtsZ-rings and arrows indicate aberrant helical FtsZ structures. Fig. S3 is related to Fig. 3A in the main text. Scale bar is 2 µm.

**Fig. S3.**

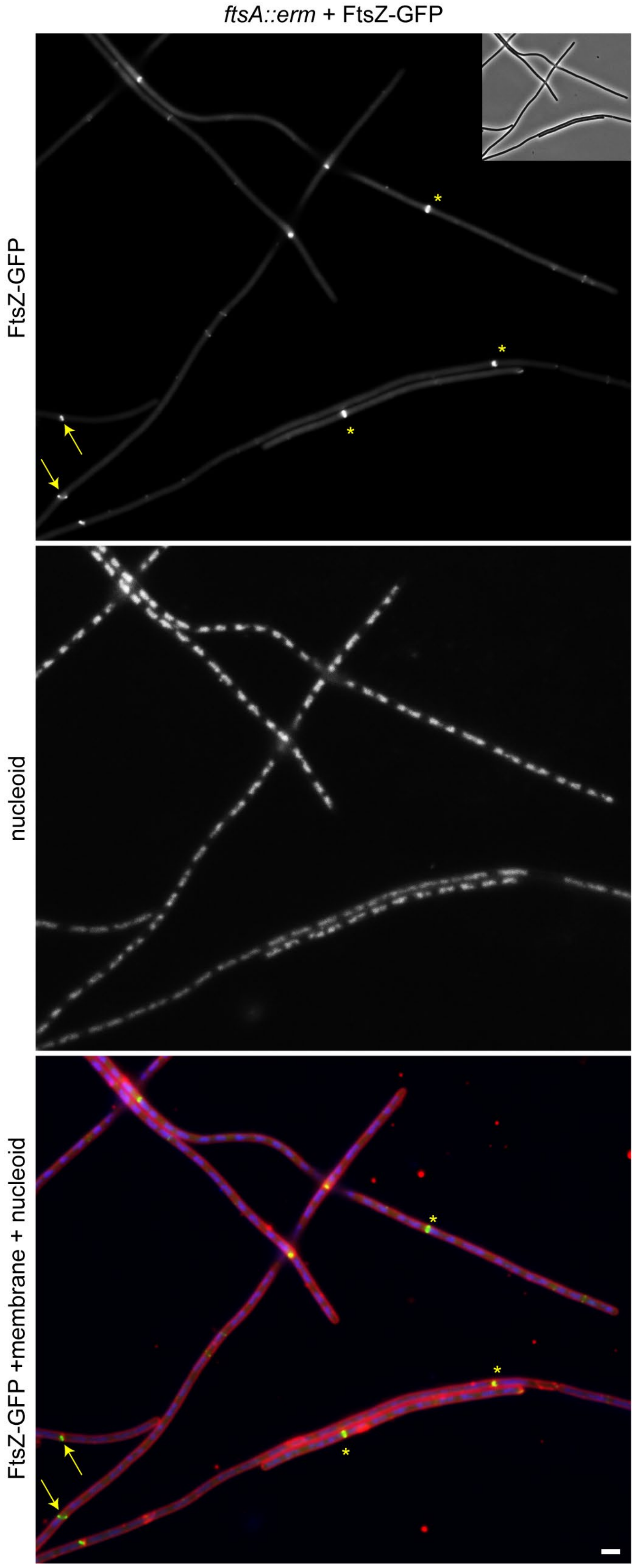


**Fig. S3. FtsZ and nucleoid localization in *ftsA::erm.***

Fluorescence microscopy images of exponential growing Δ*ftsA*::*erm* cells expressing an inducible ectopic FtsZ-GFP reporter (strain TNVS553). Phase contrast image is shown as inset in the upper panel. Membrane (red) and nucleoids (blue) were stained with FM5-95 and DAPI, respectively (2$^{de}$ and 3$^{de}$ panel, respectively). Asterisks indicate FtsZ-rings and arrows indicate aberrant helical FtsZ structures. Fig. S4 is related to Fig. 3C in the main text. Scale bar is 2 µm.

**Fig. S4.**

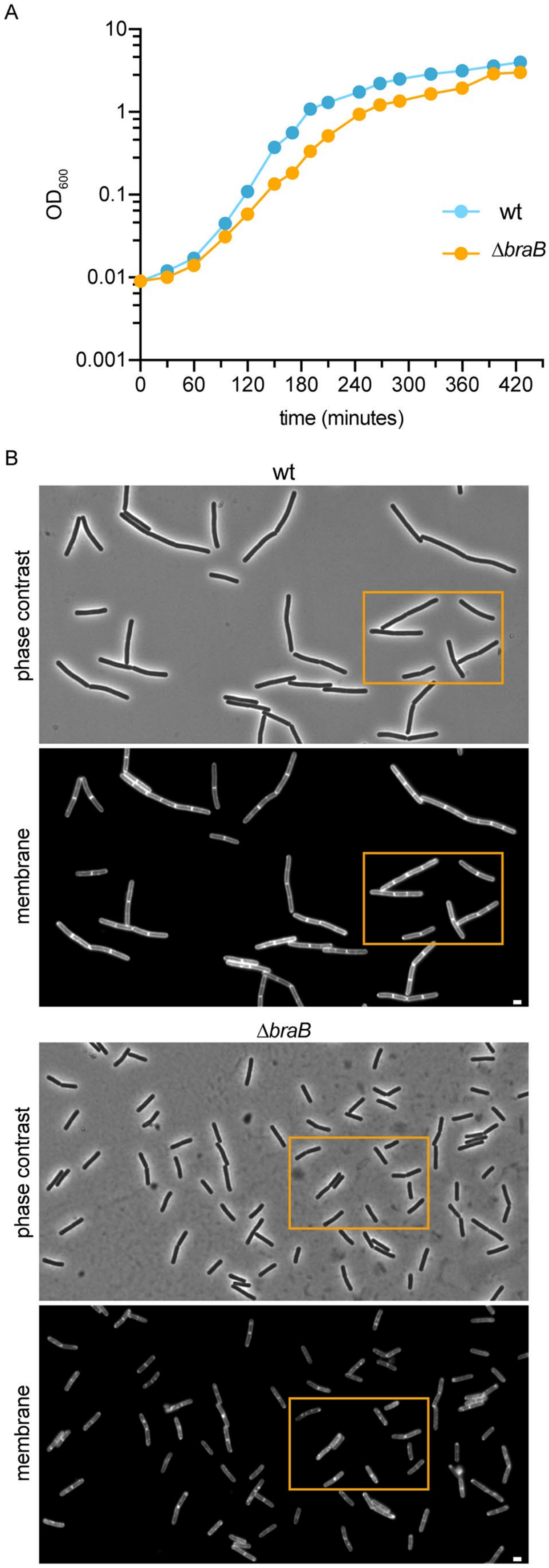


**Fig. S4. Phenotype of the *braB* deletion mutant.**

(A) Growth curves of Δ*braB* and wild type cells (wt) grown in LB medium at 37 $^{o}$C. (B) Aberrant fluorescent membrane pattern in exponential growing Δ*braB* cells (strain TNVS292) compared to wild type cells. Cells were stained with the membrane dye FM5-95. Orange boxed regions are shown in main text Fig. 8A. Scale bars are 2 µm.

**REFERENCES**


1. T. Morimoto, K. Ara, K. Ozaki, N. Ogasawara, A new simple method to introduce marker-free deletions in the *Bacillus subtilis* genome. *Genes Genet Syst* **84**, 315-318 (2009).
2. S. Ishikawa, Y. Kawai, K. Hiramatsu, M. Kuwano, N. Ogasawara, A new FtsZ-interacting protein, YlmF, complements the activity of FtsA during progression of cell division in *Bacillus subtilis*. *Mol Microbiol* **60**, 1364-1380 (2006).
3. D. R. Zeigler *et al.*, The origins of 168, W23, and other *Bacillus subtilis* legacy strains. *J Bacteriol* **190**, 6983-6995 (2008).
4. P. Gamba, J. W. Veening, N. J. Saunders, L. W. Hamoen, R. A. Daniel, Two-step assembly dynamics of the *Bacillus subtilis* divisome. *J Bacteriol* **191**, 4186-4194 (2009).
5. A. Feucht, J. Errington, *ftsZ* mutations affecting cell division frequency, placement and morphology in *Bacillus subtilis*. *Microbiology* **151**, 2053-2064 (2005).
6. B. M. Koo *et al.*, Construction and Analysis of Two Genome-Scale Deletion Libraries for *Bacillus subtilis*. *Cell Syst* **4**, 291-305 e297 (2017).
